# A graph-based learning approach to predict the effects of gene perturbations on molecular phenotypes

**DOI:** 10.64898/2026.03.20.712202

**Authors:** Yiyang Jin, Yuriy Sverchkov, Anastasiya Sushkova, Michael Ohtake, Christopher Emfinger, Mark Craven

**Affiliations:** Department of Biostatistics & Medical Informatics, University of Wisconsin, Madison, Wisconsin, U.S.A.; Institute for Immunity, Transplantation and Infection, Stanford University, Stanford, California, U.S.A.; Epic Systems, Verona, Wisconsin, U.S.A.; Barnstable Brown Diabetes Center, University of Kentucky, Lexington, Kentucky, U.S.A.

## Abstract

**Motivation:** Large-scale gene knockdown/knockout screens have been used to gain insight into a wide array of phenotypes and biological processes. However, conducting such experiments is expensive and labor-intensive. In this work, we present a general graph-based machine-learning approach that can predict the effects of gene perturbations on molecular phenotypes of interest given some measured phenotypic effects of other gene perturbations. The motivation for learning models that can predict the effects of gene perturbations is fourfold. Such models can (1) predict effects for unmeasured genes in cases in which cost or technical barriers preclude perturbing every gene, (2) prioritize unmeasured genes or sets of genes for subsequent perturbation experiments, (3) hypothesize mechanisms that underlie the relationships between the perturbed genes and their effects, and (4) generalize to other unmeasured phenotypes of interest.

**Results:** We evaluate our approach by applying it, in conjunction with four different learning methods, to learn models for four varied phenotypes. Our empirical evaluation demonstrates that the learned models (1) show relatively high levels of predictive accuracy across the four phenotypes, (2) have better predictive accuracy than several standard baselines, (3) can often learn accurate models with small training sets, (4) benefit from having multiple sources of evidence in the input representation, (5) can, in many cases, transfer their predictive value to other phenotypes.

**Data availability:** The assembled data sets and source code for this work are available at: https://github.com/Craven-Biostat-Lab/graph-molecular-phenotype-prediction

**Author summary:** One general approach for gaining insight into the genes involved in a specific biological process is to conduct an experiment in which individual genes are perturbed and the effect on the process is measured for each perturbation. Large-scale experiments of this type have provided important biological insights, but they are often expensive and labor-intensive to perform. As a result, it is not always feasible to measure the effects of perturbing every gene. In this article, we present a machine-learning approach to predicting the effects of gene perturbations using available experimental data and biological network information. Our method can estimate the effects of genes that have not yet been experimentally measured, helping researchers identify promising genes to study next. In addition, the models can suggest hypotheses about the molecular interactions that link genes to the biological process of interest. Approaches like this may help guide experimental studies and accelerate the discovery of gene–phenotype relationships.

## Introduction

Genetic perturbation methods can reveal causal relationships between genetic elements and molecular or cellular phenotypes of interest. More specifically, methods for either knocking out or knocking down genes, including CRISPR gene editing and RNA interference, can identify the genes and pathways involved in such phenotypes. High-throughput gene-perturbation screens have been used to uncover the genes involved in a broad range of phenotypes including cell morphology (1), cholesterol metabolism (2), host-pathogen interactions (3), and mitochondrial metabolism (4) among many others.

Here, we present a general, graph-based machine learning approach developed to predict the effects of gene perturbations on molecular phenotypes of interest. The motivation for learning models that can predict the effects of gene perturbations is fourfold. Such models can (1) predict effects for unmeasured genes in cases in which cost or technical barriers preclude perturbing every gene, (2) prioritize unmeasured genes or sets of genes for subsequent perturbation experiments, (3) hypothesize mechanisms that underlie the relationships between the perturbed genes and their effects, and (4) generalize to other unmeasured phenotypes of interest.

Several methods have been developed to predict scalar phenotypic changes from gene perturbations. Among them, DeepEP (5), DeepHE (6), and EPGAT (7) are knowledge-graph-based, deep learning approaches designed to predict which human genes are essential. DeepEP and DeepHE extract low-dimensional representations of genes from a protein-protein interaction (PPI) network using node2vec (8). DeepEP integrates these representations with gene expression profiles whereas DeepHE combines them with DNA-sequence features to predict essential genes. EPGAT leverages graph attention networks to predict essential genes using a PPI network, and orthology and subcellular localization features. A notable limitation of these methods is that they are restricted to predicting a single scalar phenotype (cell survival, in the applications considered). Moreover, each learned model is able to make predictions about only one phenotype.

Another class of methods have been developed to predict the effects of genetic perturbations on gene expression. Given pre-perturbation single-cell gene expression profiles and a set of perturbed genes, these models predict the resulting transcriptomic responses. GEARS (9) incorporates a Gene Ontology (GO) graph and a gene coexpression graph with deep neural networks to predict the effects of perturbations.

Similarly, BioDSNN (10) employs a variational autoencoder to generate post-perturbation gene expression profiles, utilizing the same graphs as GEARS with the addition of Reactome pathways (11). A limitation of these approaches is that they focus solely on predicting changes to gene expression profiles and do not generalize to other phenotypes.

In contrast to all these previously developed methods, our approach generalizes to a wide range of molecular and cellular phenotypes, and can learn a single model to predict multiple phenotypes.

We evaluate our approach by applying it, in conjunction with four different learning methods, to learn models for four phenotypes. The instance labels for training and testing are derived from large-scale CRISPR screens for these phenotypes, and the feature vectors are derived from a knowledge graph populated using several publicly available databases of gene/protein interactions and function. Our empirical analysis demonstrates that our learned models (1) can predict the effects of gene perturbations left out of the training set, (2) have better predictive accuracy than several baseline approaches, (3) can predict the effects of perturbations on other phenotypes not in the training set, and (4) benefit from multiple sources of evidence in the feature representation.

## Materials and methods

In this section, we describe our method, the baseline methods we compare against, and the data sets we use in our empirical evaluation.

### Overiew

The core idea underlying our approach is that the regularities of which gene perturbations will influence a phenotype can be elicited from a knowledge graph that incorporates representations of the gene, the phenotype, and the ways in which they are related via other entities and relationships. We assume that each gene is represented by a node in the knowledge graph and each phenotype of interest *P* is indicated by one or more surrogate nodes, which we refer to as *targets*. For example, one of the phenotypes we consider is cholesterol uptake, and we represent it by a single target node corresponding to the LDLR gene. We aim to learn a model to predict whether a perturbation of a *source* gene *g* will significantly affect *P*. The model can consider attributes of *g*, attributes of *P*, and how they are related through paths in the graph. Fig 1 provides an overview of the approach.

**Fig 1.**
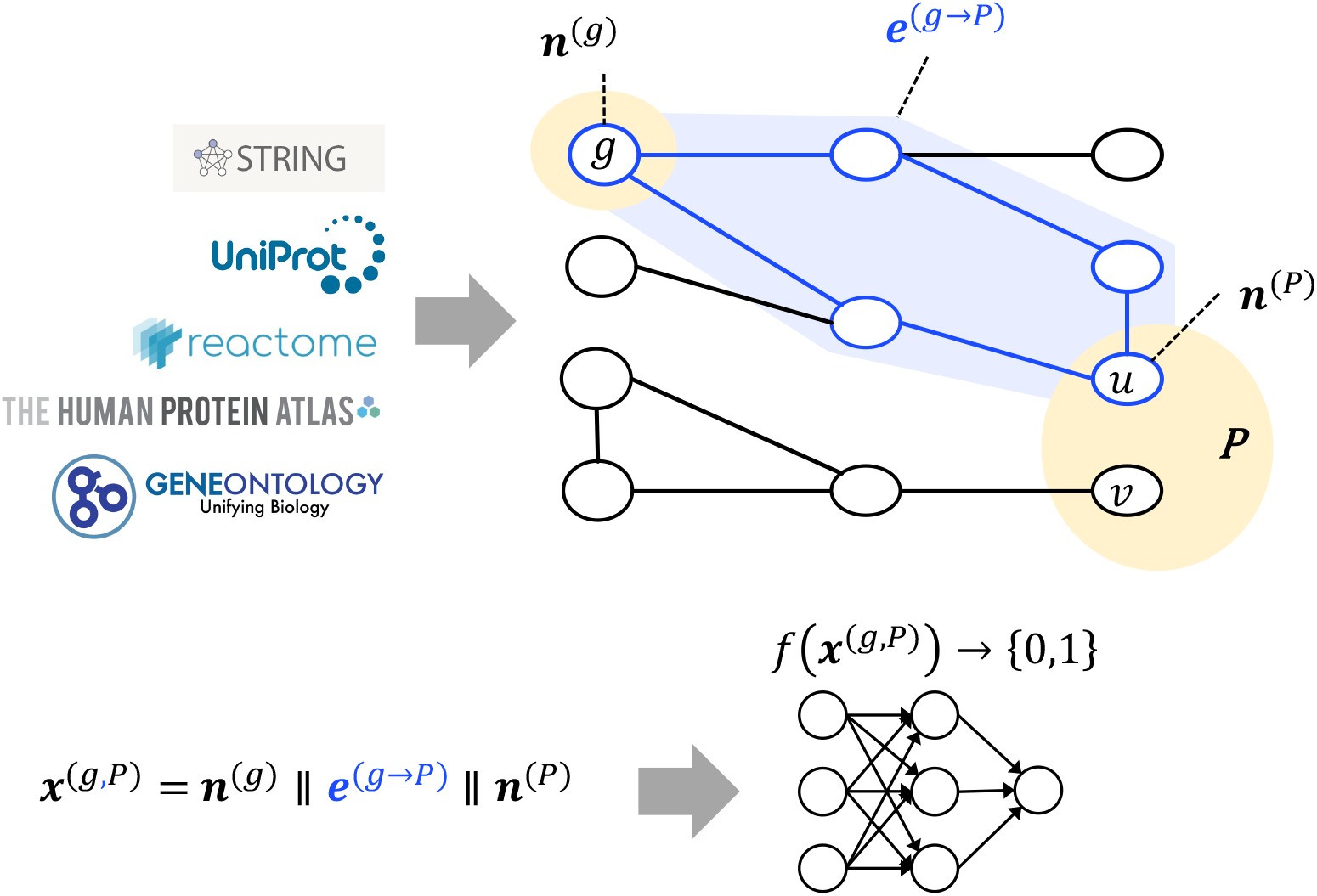
Overview of our graph-based learning approach. We populate a knowledge-graph by extracting information from multiple, public databases. A phenotype of interest *P* is represented by one or more nodes in the graph (here, *u* and *v*). To predict whether a perturbation of gene *g* will significantly affect *P*, we consider the relationship of *g* and *P* in the knowledge graph. The feature vector **x**^(*g,P*^) is constructed by concatenating a vector **n**^(*g*)^ representing node *g*, a vector **e**^(*g*→*P*^), representing the paths connecting *g* to one or more nodes in *P*, and a vector **n**^(*P*^), representing the node(s) in *P* to which *g* is connected by a path (in this example, just node *u*). A learned model *f* (*·*) takes **x**^(*g,P*)^ as input and outputs a binary value as its prediction.

The knowledge graph employed in our approach represents biomolecular entities and their interactions within a cell. For the experiments reported here, each node in the graph corresponds to a gene and the protein it encodes. More generally, the nodes could include other molecular entities such as metabolites and complexes. The edges in the graph represent interactions between the nodes. The nodes and edges both have attributes describing specific properties of the genes and interactions. The knowledge graph we assemble incorporates information from multiple, publicly available databases. The edges come from the human physical subnetwork of the STRING database (12) which contains physical and functional associations between pairs of proteins. The attributes on each edge characterizes the sources of evidence and associated confidence values that support the interaction. The attributes on the nodes describe multiple properties of each gene/protein including its subcellular localization, its abundance in various cell lines and cell types, and its functional annotations. Subcellular localization information is extracted from UniProt (13) and Reactome (11), abundance information is extracted from the Human Protein Atlas (14), and functional annotations are from the Gene Ontology (15).

The learning task we consider is to infer a model *f* (*·*) that takes a feature vector **x**^(*g,P*^) as input and outputs a binary value as its prediction, *f* (**x**^(*g,P*^)) *→ {*0, 1*}*, where 1 indicates that a perturbation of *g* will have a significant impact on *P*, and 0 indicates otherwise. The feature vector **x**^(*g,P*^) is constructed by concatenating a feature vector **n**^(*g*)^ representing attributes of source node *g*, a feature vector **e**^(*g*→*P*^) characterizing the relationship between *g* and *P* and the paths that connect them, and a feature vector **n**^(*P*^), representing attributes of the node(s) in *P* to which *g* is connected by a path.

In this setting, a model can be trained on a set of gene-phenotype pairs and then potentially be used to make predictions for genes not in the training set, phenotypes not in the training set, or both. The models can be learned using a broad range of learning algorithms that assume feature vector input representations.

### Feature Representation

As noted above, the feature vector **x**^(*g,P*^) is constructed by concatenating a vector **n**^(*g*)^ representing node *g*, a vector **n**^(*P*^), representing the nodes in *P* to which *g* is connected by paths, and a vector **e**^(*g*→*P*^) characterizing the relationship between *g* and *P* and the paths that connect them. We refer to these three constituent vectors as the *source features, target features*, and the *source-target relation features*, respectively. The features are computed using the diverse types of information represented in the knowledge graph.

The source features **n**^(*g*)^ and target features **n**^(*P*^) are computed from the same types of information and have the same dimensionality. There are three types of source and target features. The first, which we refer to as the cellular abundance features, represent each gene product’s abundance in the relevant cell line, and when available, cell type. RNA abundances for the cell line and cell type and protein abundance for the cell type are all extracted from the Human Protein Atlas.

The second set of source and target features are the subcellular localization features. We use 41 binary features to represent a hierarchical taxonomy of subcellular compartments. These features are instantiated for each gene by extracting localization annotations from both UniProt and Reactome.

The third set of source and target features provides an embedded representation of the Gene Ontology (GO) annotations for each gene. We adopt the same approach used in GEARS (9) to construct a graph representing gene-gene functional relationships, and then infer an embedded representation of each node in this graph. Let *N*_*u*_ be the set of GO annotations for gene *u*. The Jaccard index between a pair of genes *u, v* can be calculated by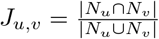, measuring the fraction of shared annotations between two genes. A GO graph is constructed by selecting the top *k* genes with highest values of *J*_*u,v*_ for each gene *u*, and adding a directed edge from *u* to each of these genes. We then apply node2vec (8; 16) to the GO graph and represent each gene by a 64-dimensional vector resulting from the learned node2vec embedding. We add to this representation a binary feature indicating the gene’s existence in the GO graph, and in cases in which a gene is not present in the graph (due to an absence of GO annotations), the 64-dimensional vector is set to all zero values.

When a phenotype is associated with multiple target nodes, the target feature vector is determined by aggregating the vectors derived for the individual target nodes. The cellular abundance and the GO features are aggregated by taking element-wise averages across the target nodes. The subcellular localization features are aggregated by taking the element-wise OR of the binary values in these features.

The source-target relation features are intended to represent various aspects of how the source gene node and the target nodes are related in the knowledge graph. One subset of these features captures the level of evidence supporting a path between the source node and a target node. Specifically, for a path of length *k*, we extract an *n*-gram of length *k* that itemizes the strongest level of evidence supporting each interaction in the path. Interactions in the STRING physical network are labeled using three evidence categories: those annotated with ‘E’ are collected from biochemical, biophysical and genetic interaction experiments, those annotated with ‘D’ are curated from pathway databases, and those annotated with ‘T’ are obtained by performing statistical co-occurrence analysis across the scientific literature. As an example, for a path consisting of three interactions with the strongest level of evidence supporting each being experimental, database, and experimental, respectively, we would extract the *n*-gram ‘EDE’. The **e**^(*g*→*P*^) vector includes an element for each such *n*-gram up to a maximum length. These features are set to counts that tally the *n*-gram occurrences for all paths between the source and a target node up to a maximum length, and then normalized to [0,1] using min-max scaling.

Each interaction in STRING is also associated with a “combined score” indicating confidence in the interaction given all sources of evidence. We compute a feature representing the product of these scores for the most confident path. Denoting a path by its sequence of nodes (*u*_1_, … *u*_*k*_), its confidence is computed as 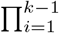score(*u*_*i*_, *u*_*i*+1_) where score(*u*_*i*_, *u*_*i*+1_) indicates the STRING combined score for the interaction between *u*_*i*_ and *u*_*i*+1_.

We also calculate a set of features that are based on the topology of paths in the graph. We include features that provide a thermometer encoding of the shortest path length, and features that record the number of paths, and an indicator as to whether there is a path between the source node and a target node. Another feature is based on the degree of the nodes on each path. We compute 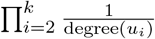 for each path (*u*_1_, … *u*_*k*_) and take the maximum as the feature value.

We also consider how a source relates to the target nodes in the graph by running diffusion processes. Specifically, we apply random walk with restart (RWR) to calculate a set of diffusion scores for each node. The scores are calculated by

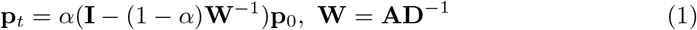

where **p**_*t*_ is a vector for post-diffusion scores in which each element corresponds to the diffusion score at time step *t* of each node in the graph, **A** denotes the adjacency matrix of the graph, **D** denotes the diagonal degree matrix of the graph, **p**_0_ is the pre-diffusion score vector, and *α* is a hyperparameter indicating the restart probability. In our approach, **p**_0_ is a binary vector in which elements corresponding to target nodes are 1 and other elements are 0 to realize the process of diffusion from the targets. We include three features representing the diffusion scores calculated by RWR with *α* set to 0.2, 0.4, and 0.6.

The final set of source-target relation features measures the similarity between the subcellular localization, GO, and the cellular abundance features for the source and the targets. Specifically, there are two features which represent the cosine similarity between the source and the target for each of the subcellular localization and GO feature groups. There is also a set of features representing the difference and the absolute difference between the source and target cellular abundance features. When a phenotype is represented by multiple target proteins, the resulting source-phenotype similarity is an aggregation (largest similarity or mean similarity) between the source and each of the targets.

### Data sets

We focus on four phenotypes of interest in our empirical analysis: cholesterol homeostasis, cholesterol uptake, influenza A virus replication, and mitochondrial protein abundance. We compile published experimental results from genome-scale CRISPR screens available in the literature, select target proteins for each phenotype, and construct data sets tailored for machine learning methods. An instance in one of our data sets is defined as a source-phenotype pair, where the source is a perturbed gene. We only consider instances for which the perturbed gene exists in the knowledge graph. Table 1 summarizes the four data sets we use in this study.

**Table 1.** Detailed description of data sets for each phenotype.

| Phenotype | Targets | # Perturbed genes | # Instances | # Positives | Cell type |
| --- | --- | --- | --- | --- | --- |
| Cholesterol homeostasis | SREBP2 | 18,224 | 18,224 | 191 | HeLa |
| Cholesterol uptake | LDLR | 17,768 | 17,768 | 473 | HepG2 |
| Influenza A virus replication | 8 host proteins w/ virus PPIs | 17,856 | 17,856 | 196 | A549 |
| Mitochondrial protein abundance | 57 mitochondrial proteins | 115 | 6,555 | 546 | HAP1 |

For the cholesterol homeostasis phenotype, we use data from two genome-wide CRISPR screens that were conducted in HeLa cells using an SREBP2-dependent transcriptional reporter (17), and by measuring plasmid membrane LDL levels mediated by SREBP pathway activation (18). We select SREBP2 as the target node for this phenotype. For the cholesterol uptake phenotype, we use data from a genome-wide screen in which the reporter was fluorescently labeled LDL-C bound to LDL receptors (LDLR) (19). We select LDLR as the target node for this phenotype. For the influenza A virus replication phenotype, we use data from three CRISPR screens that were conducted in A549 cells to identify essential host factors (20; 21; 22). Consulting the STRING database, we select eight human proteins that directly interact with influenza A proteins as the targets for this phenotype.

The set of instances for each data set is specified by pooling the genes screened across the studies. To determine which instances have positive labels, we use the following procedure. All the studies mentioned above identified genes that had a significant effect on the phenotype using MaGeCK analysis (23). We apply a uniform criterion to classify instances as positive or negative across these phenotypes and studies. Specifically, we label an instance as positive if the perturbed gene was identified as affecting the phenotype in any of the associated screens at a false discovery rate (FDR) of less than 0.1 in the reported MaGeCK analysis. Thus, when we have multiple studies per phenotype, we pool the positives identified across the studies, based on the premise that a lack of overlap in the identified positives by the studies is primarily due to false negatives rather than false positives (24). All other instances are designated as negatives.

For the mitochondrial protein abundance phenotype, we derive the data from a study that knocked out 115 genes whose products localize to mitochondria and then measured the resulting changes in abundances of 57 mitochondrial proteins (4). Each instance in this data set corresponds to a knocked out gene (the source) and one of the measured proteins (the target). We label an instance as positive if the associated *q*-value (as reported in the study (4)) is below 0.05. All other instances are labeled as negatives.

### Model Training

In this study, we apply four machine-learning algorithms to learn models to predict the effects of gene perturbations: elastic net logistic regression (25), random forest (26), XGBoost (27), and neural networks (NN). We set the maximum of number of iterations to 5000 for elastic net, and the number of trees to 1,000 for random forest. Due to the imbalance of our data sets, we use balanced class weights during training of elastic net logistic regression, random forest, and XGBoost.

The neural networks are fully connected between layers, and consist of one input layer, either one or two hidden layers, and one output unit. The hidden units use either the rectified linear unit (ReLU) activation function or the sigmoid activation function. After each hidden layer, a dropout layer is used with a dropout rate 0.3. The output unit uses a sigmoid activation function, and the loss function is binary cross-entropy loss.

A set of hyperparameters for each alogrithm are tuned using internal 5-fold stratified cross-validation within each training set. The hyperparameters are tuned using a grid search over a manually-defined parameter space with the default value of the hyperparameter covered. For elastic net logistic regression, we tune the inverse of regularization strength (C), and the elastic net mixing parameter L1. For random forest, we tune the maximum tree depth, minimum number of samples for internal node splitting, and minimum number of samples for a leaf node. For XGBoost, we tune the learning rate, maximum tree depth, minimum sum of instance weight for a child, subsampling ratio of training set, subsampling ratio of features for each tree, and L1 regulariztion on weights. For neural networks, we tune batch size, learning rate, number of training epochs, size of the hidden layer, number of the hidden layers, and the activation function used by the hidden layers. We allow the neural networks to have either one or two hidden layers, with all units using either a sigmoid or ReLU activation function. For two-layer neural networks, we set the size of the second hidden layer to be a quarter of the first hidden layer, and the first hidden layer has the same size range as the hidden layer in single-layer neural networks.

### Baseline Methods

We compare the predictive acccuracy of our learned models with two types of baseline methods: one based on path lengths, and the other based on diffusion processes. Both of these baselines are aimed at quantifying how “close” a source node is to a target node for the given phenotype. The first baseline is simply the length of the shortest path between the source node and the nearest target node in the knowledge graph. The second group of baselines we consider consists of several diffusion-process methods. Starting from a designated set of nodes, these methods propagate information through the edges to nearby nodes in an iterative manner for a fixed number of steps or until convergence (28). Each node in the graph ends up with a score after the diffusion process, and these scores can be used to rank source nodes according to their predicted effect on target nodes.

We evaluate three diffusion kernels: random walk (RW), random walk with restart (RWR), and the heat kernel. We calculate RWR scores as we do for the RWR-based features described in the Feature Representation section. We calculate RW diffusion scores for nodes in the knowledge graph by

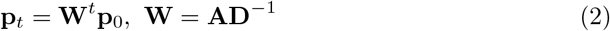

and heat diffusion scores by:

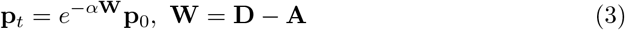

where **p**_*t*_, **p**_0_ are post and pre-diffusion score vectors, **A** and **D** are the adjacency and diagonal degree matrix of the graph, *α* is a hyperparameter, and *t* is the number of propagation steps.

For all three diffusion methods, we apply two different approaches that vary in how the pre-diffusion vectors **p**_0_ are initialized. The first, which we refer to as *target diffusion*, sets elements in **p**_0_ that correspond to target nodes as 1 and all other elements to 0. In effect, the diffusion process starts only from the target nodes. In the second approach, which we refer to as *positive diffusion*, the elements of **p**_0_ that correspond to the positive instances in the training set, in addition to the targets, are initialized to 1, and all other elements are set to 0. We can think of this diffusion process as starting from known positive source genes in addition to the targets for the phenotype.

## Results

In this section, we present a set of experiments designed to evaluate the predictive accuracy and generality of our approach.

### Learned models show high predictive accuracy

We first test the ability of the learned models to predict the effects of held-aside gene perturbations on the phenotypes of interest by performing 5-fold cross-validation within each data set. For the phenotypes cholesterol homeostasis, cholesterol uptake, and influenza A virus replication, the cross validation is done using stratified sampling so that each fold contains approximately the same number of positive and negative instances. Each training set also includes a special instance for each target node in which the target itself is treated as the source node, given that we know that the target nodes are always important for the phenotype. For the mitochondrial protein abundance data set, we divide the 115 knocked-out genes into five folds so that each fold contains the same number of source genes. We construct ROC curves by pooling the test-set predictions across the folds for each data set.

We learn models by applying four machine learning methods – elastic net logistic regression, random forest, XGBoost, and neural networks – to each data set. For all four phenotypes, we use internal 5-fold cross-validation within each training set to select the hyperparameters for that training set.

Fig 2 presents the ROC curves for all four learning methods across the four phenotypes. With an average AUROC of 0.72, the learned models demonstrate strong predictive performance across different phenotype scenarios. Moreover, the varied learning methods demonstrate comparable predictive accuracy across all phenotypes. Both of these results provide evidence of the generality of our approach.

**Fig 2.**
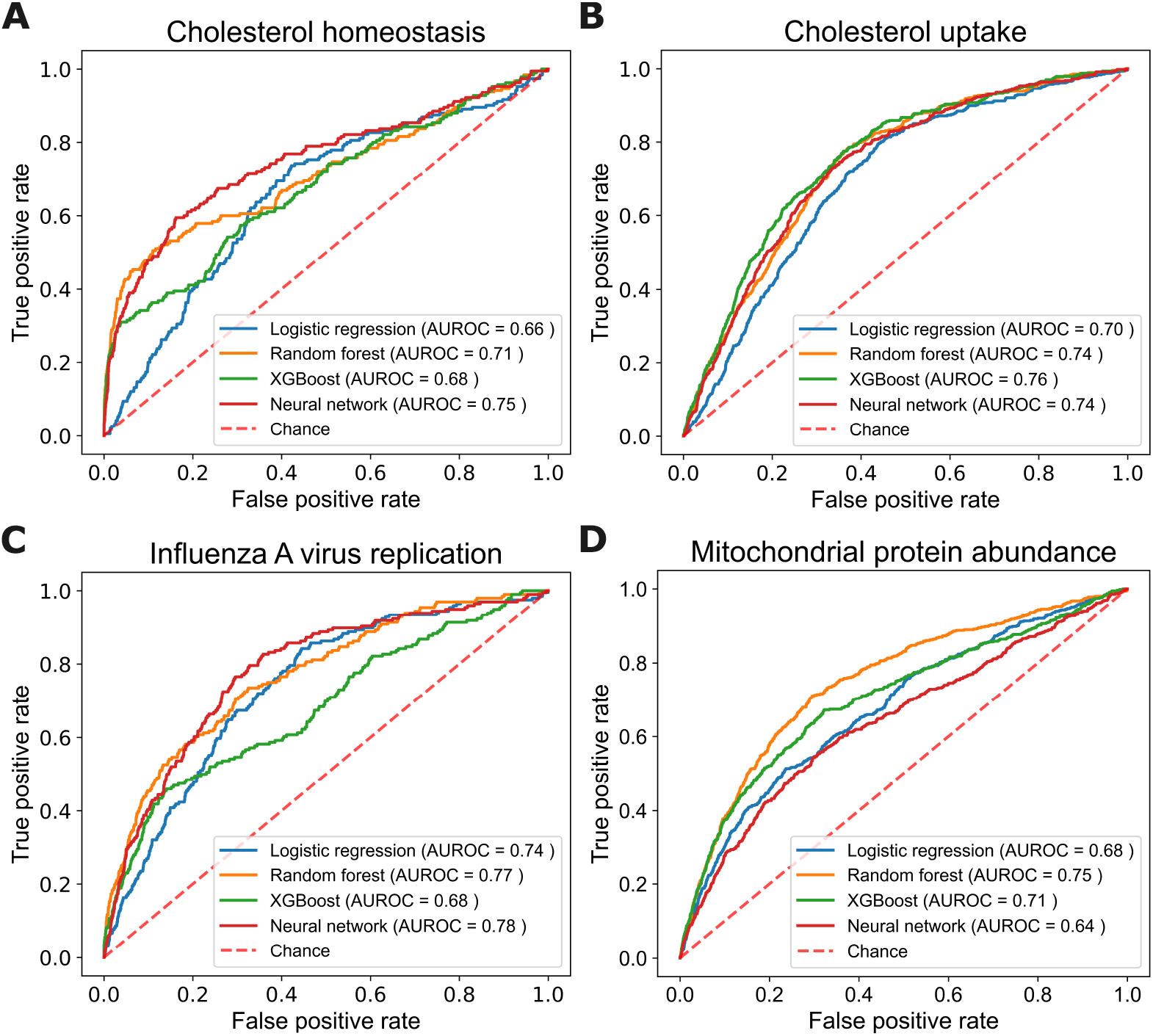
Predictive accuracy of the four learning algorithms for the four phentoypes. Each panel shows the ROC curves and AUROC values for each learning method on the given phenotype.

### Learned models have better predictive accuracy than baselines

To further benchmark our approach’s predictive accuracy, we compute learning curves and compare learned models to the baseline methods described in the Baseline Methods section. For each instance, we define its shortest-path-length baseline score as the shortest path length from the source to the nearest target. For the diffusion baselines, we define an instance’s score as the post-diffusion score of the instance’s source node. We compute target diffusion and positive diffusion scores using a range of hyperparameters: for the RW kernel, we set *t* to 10, 20, and 30; for the RWR kernel, we set *α* to 0.2, 0.4, and 0.6; and for the heat kernel, we set *α* to 0.1, 0.2, and 0.3. For each baseline method we report only the *best* result across the hyperparameter settings.

For the learning methods, we adopt the same hyperparameter tuning strategy as in the previous section, conducting internal 5-fold cross-validation for each training set size for each phenotype. To analyze how predictive accuracy changes as a function of the amount of training data, we generate training sets of varying sizes by randomly sampling instances from each complete training set without stratification. We plot learning curves for the learning methods and for the positive diffusion baseline methods.

Fig 3 shows the learning curves for the baseline methods and our machine-learning approach. The *x*-axis shows the fraction of the available training set used and the *y*-axis shows the pooled AUROC from cross validation. Because the target diffusion scores and shortest path length baseline do not make use of a training set, their AUROC values remain constant across the *x*-axis. We observe several interesting results in these plots. First, the learning methods generally show increasing trends in AUROC values as the training set size grows, sometimes reaching near-maximum AUROC even with a relatively small subset of the available data.

**Fig 3.**
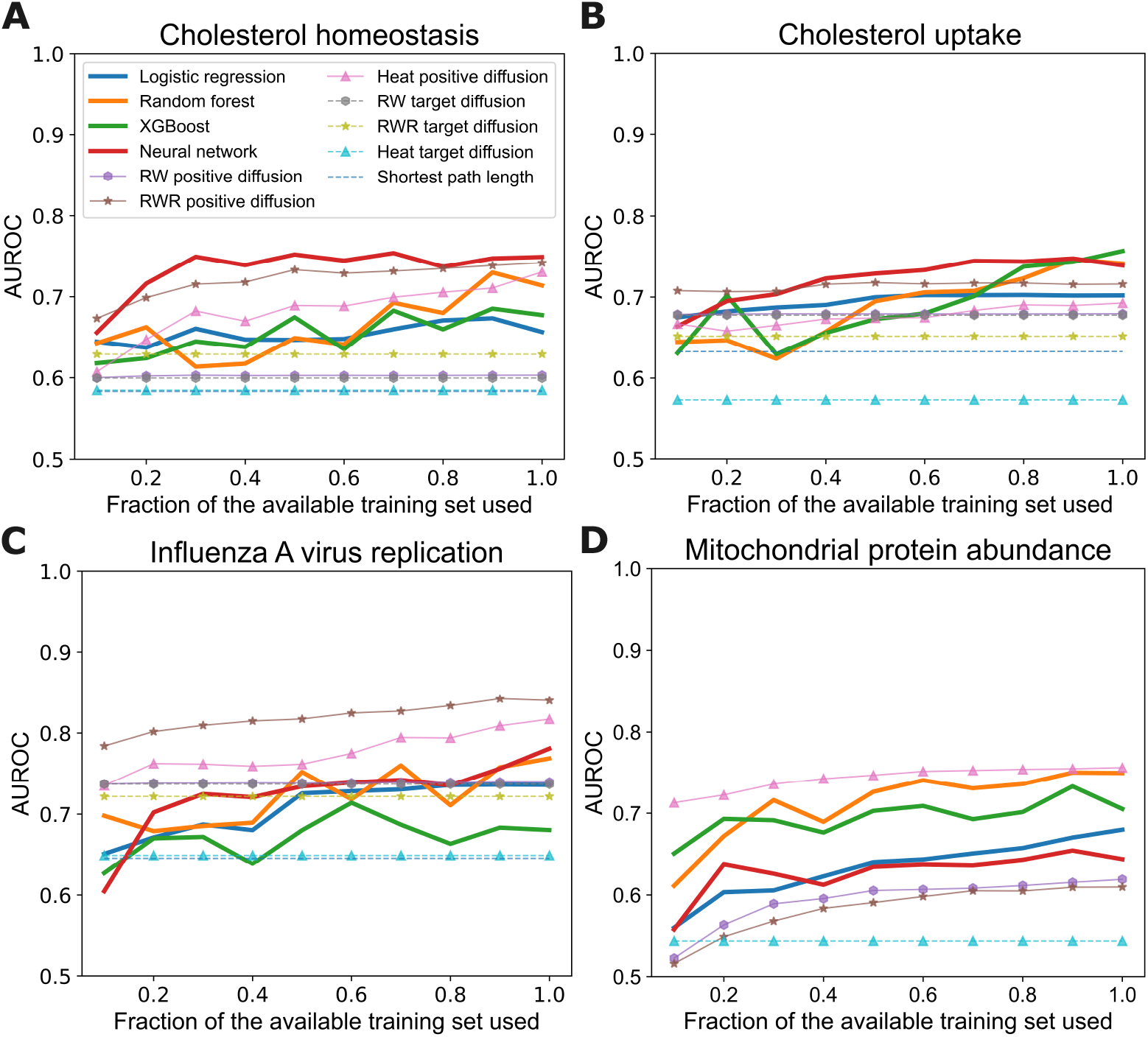
Learning curves for baseline methods and the learning methods. Each panel shows the results for one phenotype. The *x*-axis shows the fraction of the available training set used and *y*-axis shows the resulting AUROC values.

Second, the learning methods exceed the predictive performance of the shortest path length and the target diffusion baselines for all phenotypes. Recall that the target diffusion approaches start the diffusion process only from the target nodes.

Third, the highest predictive accuracy is achieved by a learned model in three out of the four phenotypes. However, the RWR positive diffusion and the heat positive diffusion baselines demonstrate competitive predictive accuracy for most phenotypes and superior accuracy for the influenza A virus replication phenotype. Recall that the positive diffusion approaches make use of a training set by starting the diffusion process from positive instances in the training set in addition to the target nodes. The relatively high predictive accuracy of these two baselines suggests that the genes that are involved in these phenotypes may be in close proximity in the knowledge graph, and thus especially amenable to prediction via diffusion.

We note again that the diffusion baselines are advantaged in this analysis because we report the best predictive performance over a range of hyperparameter settings for each. Moreover, we note that a significant limitation of the the positive diffusion methods relative to our approach is that they cannot generalize to other phenotypes without having training sets specifically for those phenotypes. In contrast, our approach can be applied to make predictions for the effects of perturbations on phenotypes that were not represented in the training set. We explore this setting in detail in a subsequent section. Overall, we conclude that our learning approach provides a way to predict the impact of gene perturbations on a phenotype with a high degree of accuracy and generality.

### Models benefit from multiple sources of evidence

The feature vector comprises features representing properties of the source node **n**^(*g*)^ and target node(s) **n**^(*P*^), along with source-target relation (ST-relation) features **e**^(*g*→*P*^) characterizing the relationship between *g* and *P* and the paths that connect them.

These features are derived from diverse data types including protein-protein interactions, cellular abundances, subcellular localizations, and GO annotations. In this section, we evaluate the predictive value of various aspects of this feature representation.

We decompose **x**^(*g,P*^) into distinct feature groups and evaluate the predictive accuracy of models learned from each group separately. We consider two types of feature groupings. First, we consider what we refer to as the three *subgraph* feature groups, which are the source node features **n**^(*g*)^, the target node features **n**^(*P*^), and the source-target relationship (ST-relation) features **e**^(*g*→*P*^). Second, we consider what we refer to as the *evidence-type* feature groups which are based on the type of evidence from which they were derived. These groups are the cellular abundance features, subcellular localization features, GO annotation features, and the features that are based on the PPI network, including those that characterize paths between the source and target nodes and the diffusion features. To systematically evaluate the predictive power of these feature groups, we learn models for each group using the same 5-fold cross-validation method for all four phenotypes, as before.

Fig 4 shows the predictive accuracy of models learned from individual subgraph feature groups. For comparison, the figure also shows the accuracy of models trained on all of the features. Bars are not shown for the cases in which the resulting AUROC *≤* 0.5. Error bars represent the AUROC variation across the 5-fold cross-validation.

**Fig 4.**
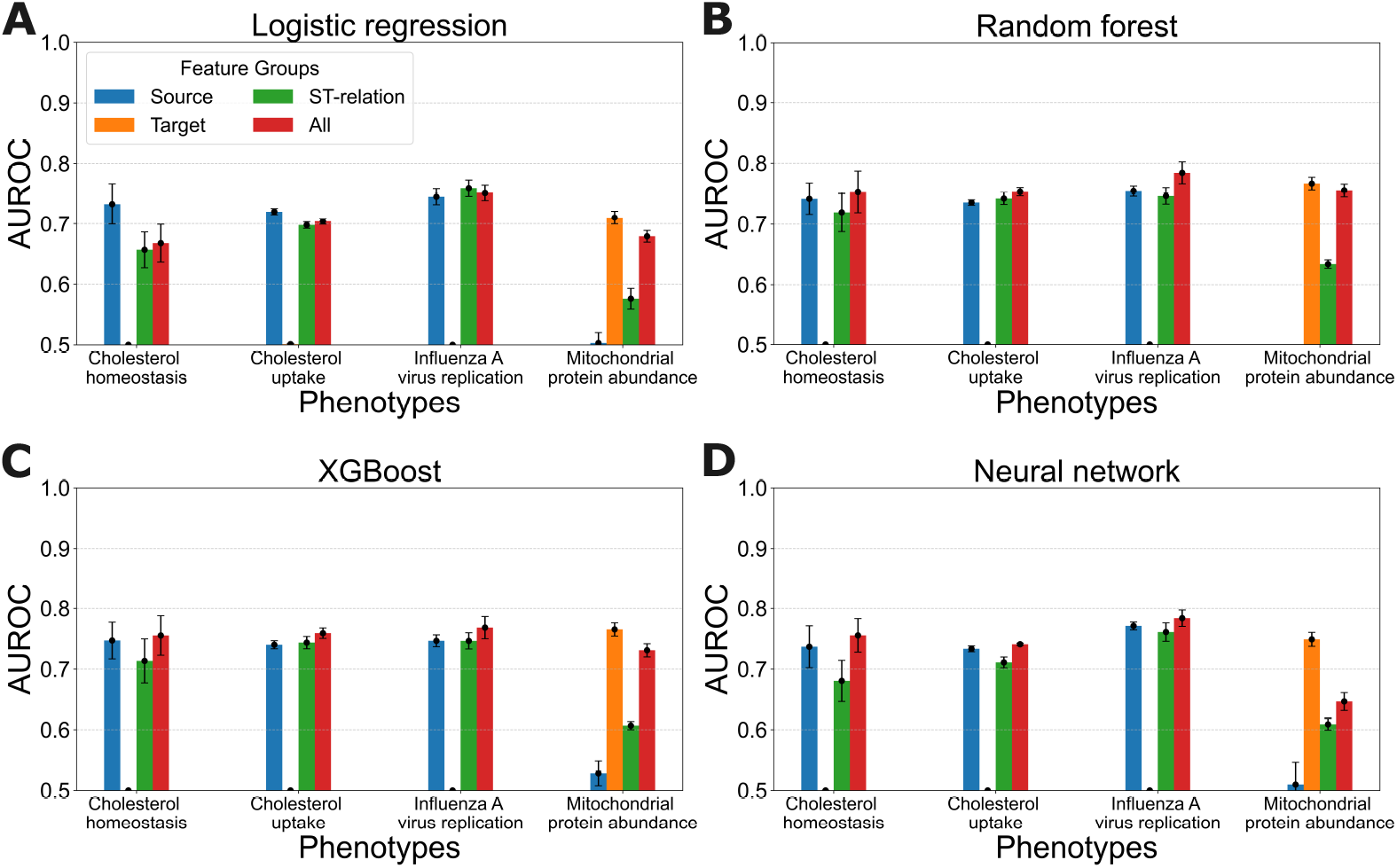
Predictive accuracy of learned models trained on individual subgraph feature groups versus models trained on all features. Each panel shows the results for a learning algorithm applied to all four phenotypes.

We observe that models trained with all features either achieve the highest or second highest mean AUROC values in all cases, indicating that the learning approach benefits from incorporating information about source nodes, target nodes, and the relationships between them.

We further observe that target features are highly predictive for mitochondrial protein abundance, whereas source features are more informative for the other three phenotypes. This observation aligns with the fact that the target features remain unchanged for cholesterol homeostasis, cholesterol uptake, and influenza A virus replication, making them less useful for distinguishing between instances in these phenotypes. Conversely, source features are not predictive for mitochondrial protein abundance due to the fact that many instances share the same source nodes in this data set.

Notably, the source-target relation features consistently contribute to predictive accuracy across all phenotypes, highlighting their importance in capturing interactions between genes and the phenotypes they influence. We note that this result is especially compelling because these features, and the models learned from them, do not encode any information about specific source and target nodes, and thus can potentially generalize across phenotypes. We explore this issue further in the next section.

Fig 5 presents the predictive accuracy of models learned from individual evidence-type feature groups. The figure also shows the accuracy of models trained on all of the features as a baseline for comparison. From these results we observe that, for most learning methods and phenotypes, using all of the features provides the best predictive accuracy. Moreover, the relative predictive value of individual feature groups varies across the phenotypes. For example, whereas subcellular localization is the most predictive individual feature group for cholesterol homeostasis, GO annotations are the most predictive group for mitochondrial protein abundances. We also observe that every evidence-type feature group has predictive value.

**Fig 5.**
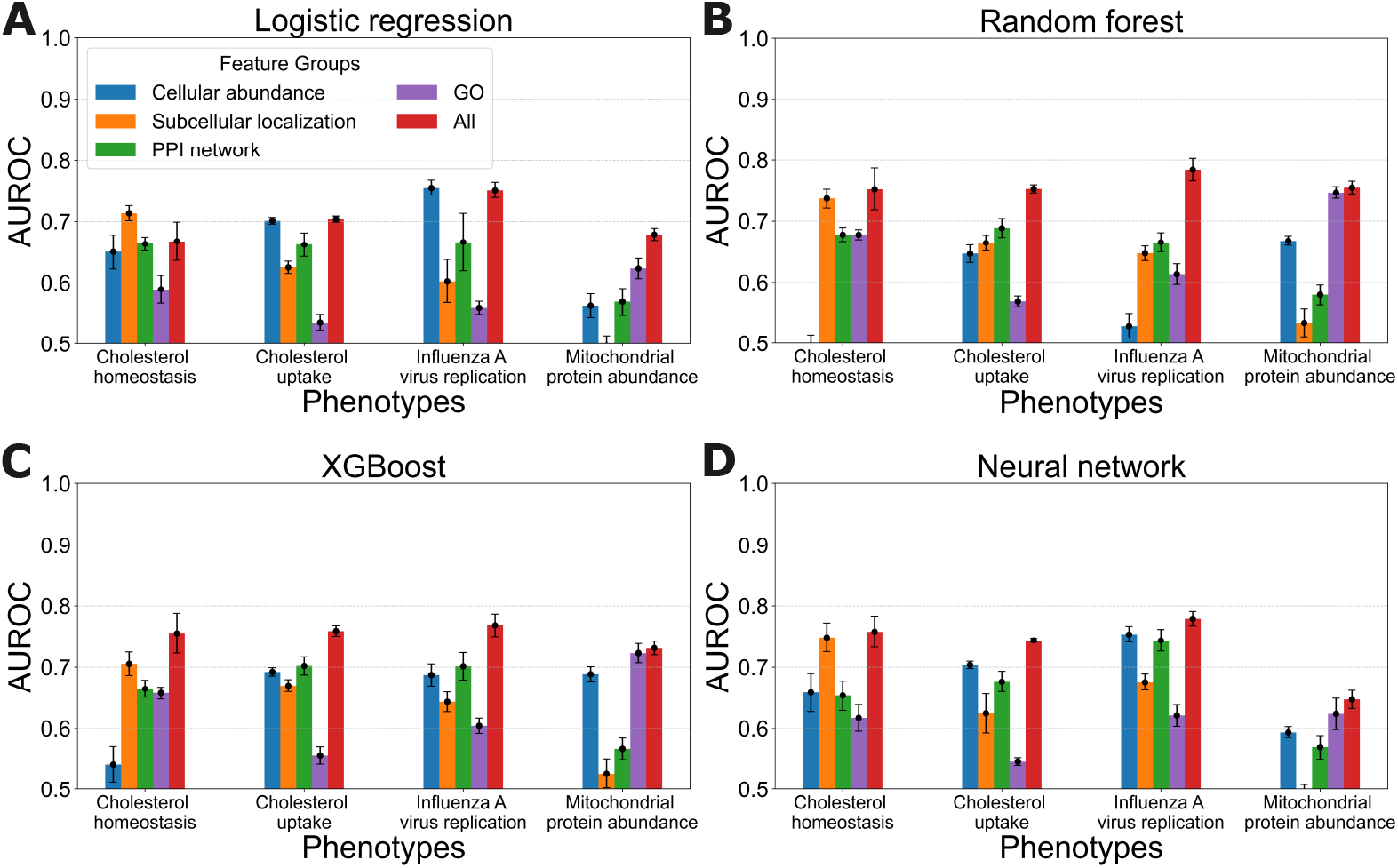
Predictive accuracy of learned models trained on individual evidence-type feature groups versus models trained on all features. Each panel shows the results for a learning algorithm applied to all four phenotypes.

In summary, the results presented in this section illustrate that our learning approach gains predictive value and generality by incorporating features that are derived from multiple sources of evidence and that represent properties of the source node, the target nodes, and the relationships between them.

### Models can predict the effects of perturbations on other phenotypes

One advantage of our approach is that a model can be trained on one phenotype and then applied to make predictions for a different phenotype. In this section, we evaluate the effectiveness of our approach in this setting by training and testing on different pairs of the phenotypes considered in the previous experiments. We do this using models that are trained using nearly the full set of features as well as models trained using only the ST-relation features. In both cases, we exclude a few of the features. which are not available across all phenotypes (e.g. some of the cellular abundance features) or are phenotype-specific (e.g. the diffusion features).

To learn models for each phenotype, we perform internal 5-fold cross-validation with stratification to select hyperparameters, and then train on the entire data set for the phenotype using the selected hyperparameters. We plot the AUROC for each learned model when it is tested on each of the other phenotypes. As a baseline for comparison, we also report within-phenotype predictive accuracy as determined in the 5-fold cross-validation experiments reported earlier.

Fig 6 shows the predictive accuracy of the trained models when tested on each other phenotype. The models generally exhibit significant predictive value in this transfer learning setting. The exception is that the mitochondrial protein abundance phenotype does not lend itself to transfer with the other phenotypes. The models trained on other phenotypes do not predict well on mitochondrial protein abundance, and vice versa.

**Fig 6.**
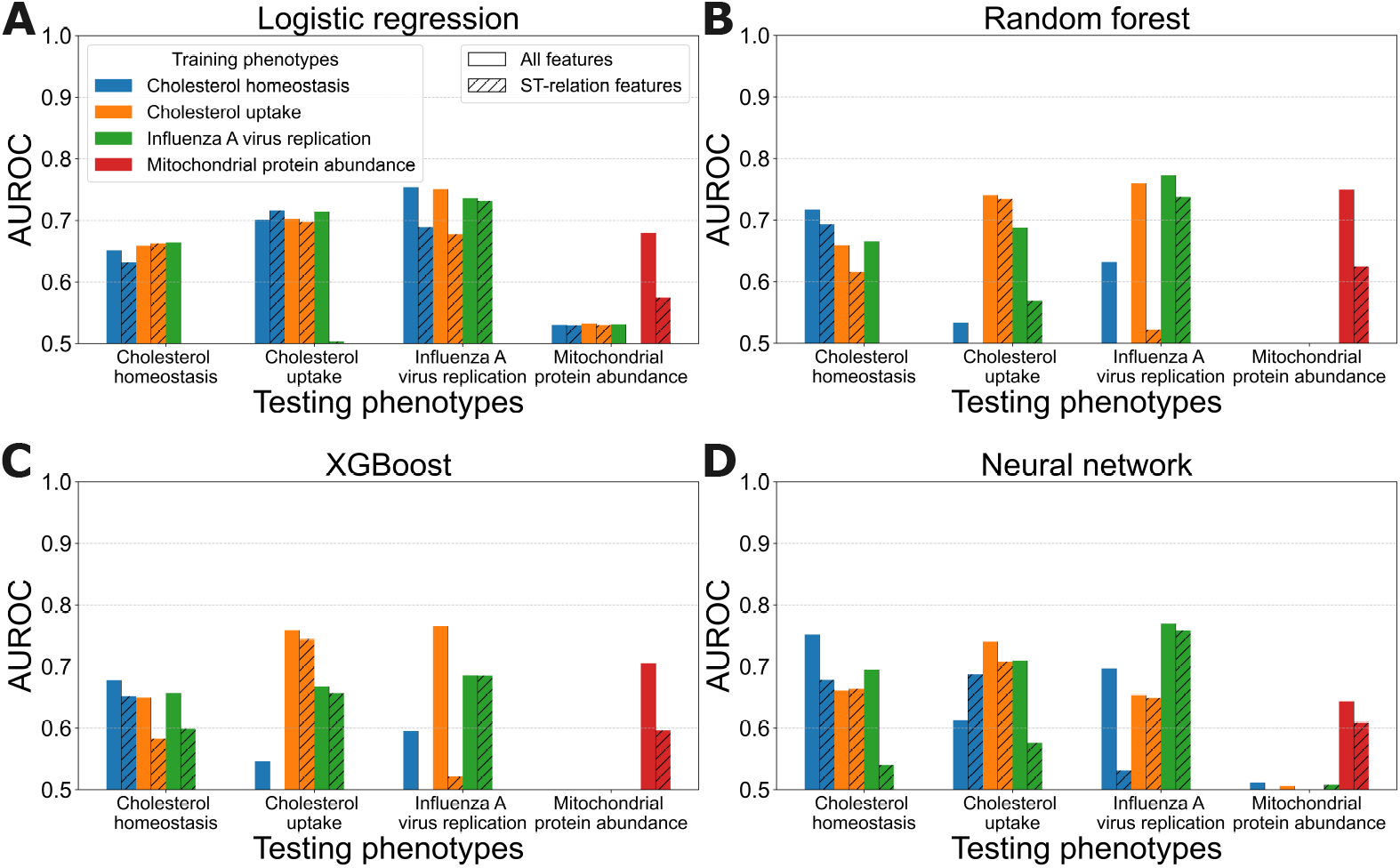
Predictive accuracy of learned models trained and tested on different phenotypes. Each panel shows the results for one learning method. Test-set phenotypes are grouped along the *x*-axis, training-set phenotypes are indicated by colored bars, and the feature sets used are distinguished by bar fill styles: solid for all features and hatched for ST-relation features

This result is perhaps due to fact that the models for this phenotype rely heavily on the target feature groups, as shown in the previous section, whereas the other phenotypes do not.

Models trained using only ST-relation features also demonstrate transferable predictive ability in most cases, although their performance is generally lower than that of models trained on all features.

### Predictive accuracy is not overly sensitive to the definition of negative instances or target nodes

Two key aspects of assembling a data set for our approach are assigning class labels to instances and specifying the target nodes that represent the phenotype. In this section, we explore the sensitivity of our models to choices in how these steps are done.

When forming the data sets for three of the phenotypes (those with studies that used the MaGeCK method), we specified negative instances for all of the knocked-out genes in the study that did not meet the FDR *<* 0.1 criterion. However, this criterion might be too stringent in defining positives and too lenient in defining negatives. Here we consider data sets in which we attempt to stringently define both positive and negative instances, and leave out of the data set the instances in the excluded middle.

To explore the impact of different definitions of the negatives, we assembled new data sets for the phenotypes of cholesterol homeostasis, cholesterol uptake, and influenza A virus replication. For the cholesterol homeostasis and cholesterol uptake phenotypes, we define the negatives as those with MaGeCK reported FDR *>* 0.9. For the influenza A virus replication phenotype, two rounds of CRISPR screening were conducted. Therefore, we define negative instances as those with genes which were perturbed in the first round but failed to be included in the second round. Our definition of the positives in these data sets is not changed. With the more stringent criterion for defining negatives, we end up with 15,488 instances for cholesterol homeostasis, 683 instances for cholesterol uptake, and 16,689 for influenza A virus replication. We evaluate random forest models on these data sets using 5-fold cross-validation and compare the predictive accuracy to the original models.

Panels A-C in Fig 7 show the ROC curves of random forest models across three phenotypes with both definitions of negative instances. We observe that the ROC curves for the original negatives and the stringently defined negatives are very close for the influenza A phenotype, with a small decay in accuracy for the former in the cholesterol homeostasis phenotype and a small boost for the latter in the cholesterol uptake phenotype. We conclude that the predictive accuracy of our learned models is not especially sensitive to the specification of negative instances.

**Fig 7.**
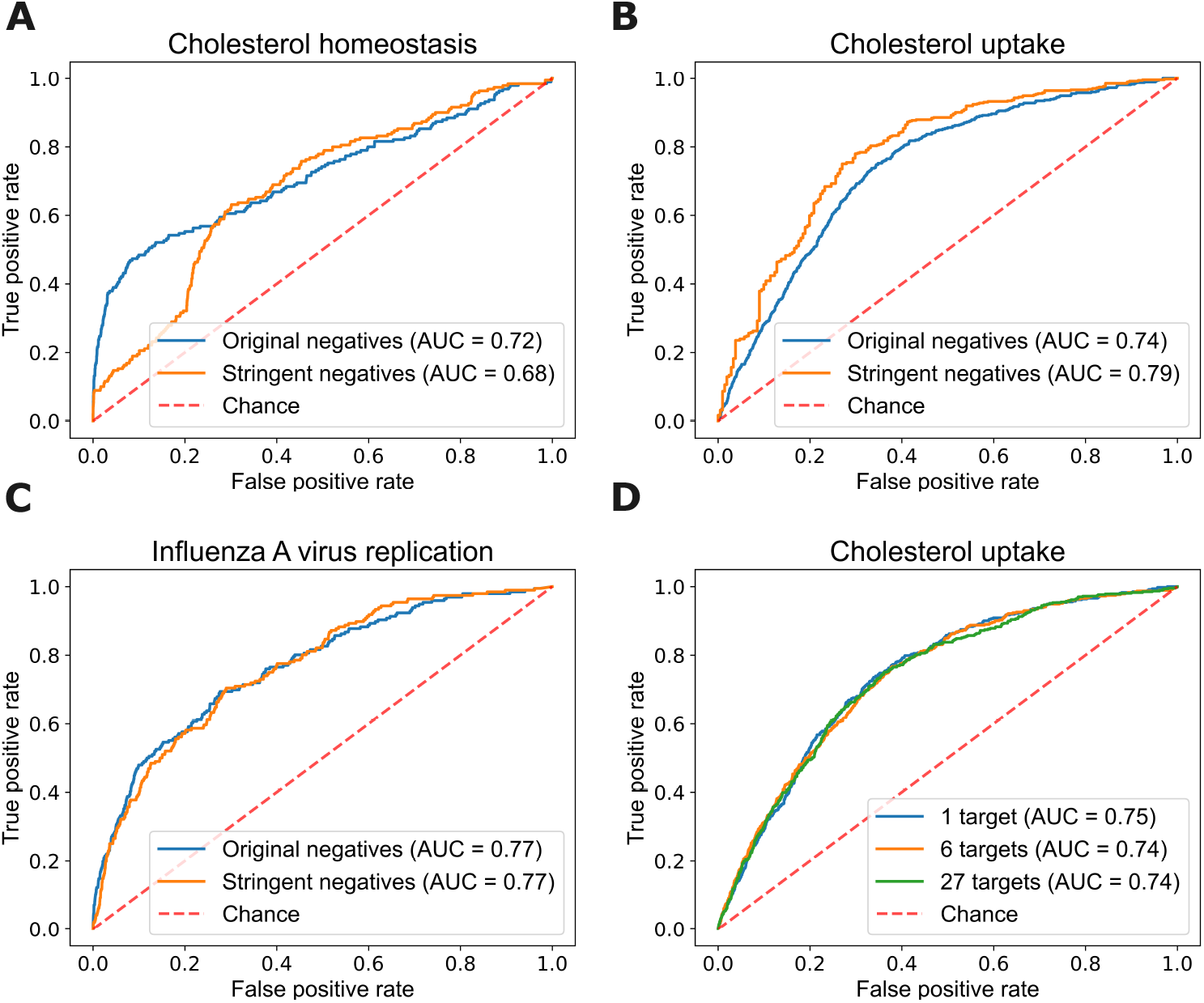
ROC curves for random forest models learned on different definitions of negative instances across three phenotypes (panels **A** to **C**), and on different target definitions of target nodes for the cholesterol uptake phenotype (panel **D**).

To explore the impact of different definitions of targets for a given phenotype, we assemble three variations of the data set for the cholesterol uptake phenotype that differ in the set of targets used to compute feature vectors. Our original representation is based on one target, LDLR. A second representation uses 27 target nodes consisting of genes annotated with two relevant Gene Ontology terms. A third, intermediate, representation uses six targets. We evaluate these different target representations by running 5-fold cross validation with the random forest method.

Panel D in Fig 7 shows the predictive accuracy of random forest models learned with these different target definitions for cholesterol uptake. We observe that the three ROC curves are nearly coincident, and thus we conclude that our approach is not especially sensitive to the exact specification of the targets, at least for the phenotype considered.

## Discussion

We have proposed a knowledge graph-based and learning method-agnostic approach that can predict the effects of gene perturbations on phenotypes of interest. The core idea underlying our approach is that the regularities of which gene perturbations will influence a phenotype can be elicited from a knowledge graph that incorporates representations of the gene, the phenotype, and the ways in which they are related via other entities and relationships in the graph. Our knowledge graph is assembled from diverse sources of evidence including protein-protein interactions, cellular abundances of gene products, protein subcellular localizations, and Gene Ontology annotations.

We conducted an empirical evaluation applying four machine-learning methods to four data sets, each of which characterizes the effects of gene knockouts on a phenotype of interest. Our evaluation demonstrated that the learned models (1) show relatively high levels of predictive accuracy across the four, diverse phenotypes, (2) have better predictive accuracy than several standard baselines, (3) can often learn accurate models with small training sets, (4) benefit from having multiple sources of evidence in the input representation, (5) can, in many cases, transfer their predictive value to other phenotypes. Additionally, we showed that our approach is not especially sensitive to how we define negative instances, or to how a phenotype is represented as a set of target nodes.

The key limitations of our approach are twofold. First, there is much room for improvement in the accuracy of the predictions the models make. We posit that this is due, in part, to noise in the genome-wide screens and their propensity to output many false negatives (24). However, we hypothesize that the predictive accuracy of the approach can be improved by augmenting the knowledge graph and by training models with more data. A second limitation of the approach is that it assumes that a phenotype can be represented by a set of surrogate target nodes in the knowledge graph.

There are several directions that we will pursue in future research to address the limitations of our approach. First, we plan to expand the types of interactions and attributes included in the knowledge graph. For example, transcriptional regulatory relationships and gene orthology information are additional types of evidence that we could include. Second, we plan to assemble additional data sets for training and evaluating models. Third, we plan to investigate extensions to our approach for handling phenotypes that do not obviously map to a small number of target nodes. One idea here is to extend the graph to incorporate process or pathway nodes. Fourth, we will investigate the utility of interpretability methods for gaining mechanistic insight into the relationships between genes and the phenotypes they influence. Fifth, we plan to adapt the approach to learn models based on graph neural networks. We believe that these extensions will advance this work towards the goal of being able to predict the effects of arbitrary gene perturbations on virtually any molecular or cellular phenotype.

## Competing Interests

No competing interest is declared.

## Author Contributions Statement

YJ, YS and MC devised the overall approach. NS and MO contributed to knowledge graph construction, data set assembly, and some of the initial experiments. YJ carried out the computational experiments. YJ and MC led the writing of the manuscript. CE contributed to the review of experimental results and the manuscript.

## Acknowledgments

We gratefully acknowledge research support from NIH/NHGRI grant U01 HG012039.

## Notes

### Competing Interest Statement

The authors have declared no competing interest.

### Summary of Updates

Figure2 revised; Figure3 revised; Figure 4 revised; Figure 5 revised; Figure 6 revised

